# Proteomic Characterization of Spodoptera frugiperda Granulovirus Occlusion Bodies

**DOI:** 10.1101/2025.06.19.660566

**Authors:** Tomás Masson, María Laura Fabre, Santiago Gómez Bergna, Matías Luis Pidre, Silvana Tongiani, Ricardo Salvador, Víctor Romanowski, María Leticia Ferrelli

## Abstract

Lepidopteran-infecting baculoviruses represent one of the most promising alternatives for pest biocontrol. In particular, Spodoptera frugiperda granulovirus (SfGV) infects the fall armyworm *Spodoptera frugiperda*, an increasingly important pest for maize and other crops around the world. SfGV has been investigated due to its capacity to enhance other baculovirus infectivity and as a biopesticide itself. Although the proteome of SfGV occlusion bodies (OB) is fundamental for their infectivity properties, we still lack a detailed understanding of its protein components and how they compare with other baculoviruses. To tackle this problem, we performed mass spectrometry-based proteomics analysis on SfGV OB obtained from infected larvae. We could detect 72 proteins included in SfGV OB and confirm the presence of two enhancins. EmPAI-based semi-quantification highlighted the presence of two granulovirus proteins, ORF101 and ORF141, shared with other GV proteomic studies. ORF101 was shown to form aggregates in insect cells. Bioinformatic analysis determined that ORF101 is an Ac5 family protein and at the same time shares striking similarities with AcMNPV P10. Prediction and comparison of OB protein structures allowed the detection of a common structural fold among 6 proteins PEP-1, PEP-2, Pep-P10, ORF007, ORF025, and Bro-f. Finally, comparison of four reported ODV proteomic studies of different GVs indicated that ORF040, ORF101, ORF141 and Bro-f are common components in GV OBs.

## Introduction

Baculoviruses are insect-specific viruses widely used around the world to control lepidopteran pests in agriculture. Their specificity, biosafety, persistence, and ease of application make them an attractive alternative to chemical insecticides or transgenic events that can trigger resistance in insect pests. *Baculoviridae* is a family of enveloped, insect viruses with a large, circular, dsDNA genome packaged in a rod-shaped nucleocapsid. Actual taxonomy classification places Baculoviruses in the order *Lefavirales*, class *Naldaviricetes*, along with other baculo-like viruses (van Oers et al., 2023). This class is not assigned to any realm, the highest official rank in virus taxonomy. Two of the six realms of the virosphere involve large dsDNA genomes (i.e. Varidnaviria or Duplodnaviria). Still, baculoviruses couldn’t be assigned to any of these due to lack of sequence homology and scarce information on the capsid proteins. Recently, based on nucleocapsid architecture revealed by cryoEM and the structures of baculovirus core genes, a new realm was proposed, termed “Telodnaviria”, for baculo-like viruses (Johnstone et al., 2024).

During their life cycle, baculoviruses produce two distinct infectious forms: budded virus (BV) and occlusion-derived virus (ODV). BVs are produced when nucleocapsids bud from the infected cells, and are responsible for dissemination of the infection in the host tissues. ODVs are produced late in infection, obtaining their envelope from the cell nucleus, and are finally embedded in a proteinaceous crystalline matrix, forming the occlusion bodies (OBs). OBs can include one to several ODVs, conferring them protection and persistence in the environment upon release from dead larvae. When OBs are ingested by a susceptible larva, OBs are dissolved in the alkaline midgut, releasing ODVs for primary infection (Rohrmann, 2019).

According to the morphology of the OB, Baculoviruses can be divided into two groups: Nucleopolyhedrovirus (NPV) and Granulovirus (GV). In terms of taxonomy, the Baculovirus family is divided into four genera: *Alphabaculovirus* (lepidopteran-specific NPV), *Betabaculovirus* (lepidopteran-specific GV), *Gammabaculovirus* (hymenopteran-specific NPV), and *Deltabaculovirus* (dipteran-specific NPV). The most widely investigated are Alpha-and Betabaculoviruses, due to their several advantages for their application as bioinsecticides to control lepidopteran pests (Harrison et al., 2018)

Due to the importance of the OB and ODV in the persistence of the infective particles in the environment and in primary infection, several studies were performed to investigate the protein composition of these particles. Up to date, nine baculoviruses were subjected to proteomic studies of their ODV or OB. These are: Autographa californica multiple nucleopolyhedrovirus (AcMNPV) (Braunagel et al., 2003), Culex nigripalpus nucleopolyhedrovirus (CuniNPV) (Perera et al., 2007), Bombyx mori Nucleopolyhedrovirus (BmNPV) (Guo et al., 2017; Liu et al., 2009), Chrysodeixis chalcites nucleopolyhedrovirus (ChchNPV) (Xu et al., 2011), Helicoverpa armigera nucleopolyhedrovirus (HaNPV) (Hou et al., 2013), Anticarsia gemmatalis multiple nucleopolyhedrovirus (AgMNPV) (Braconi et al., 2014), Mamestra brassicae nucleopolyhedrovirus (MabrNPV) (Hou et al., 2016), Pieris rapae granulovirus (PrGV) (Wang et al., 2011), Clostera anachoreta granulovirus (ClanGV) (Zhang et al., 2015), and Epinotia aporema granulovirus (EpapGV) (Masson et al., 2019).

Spodoptera frugiperda granulovirus (binomial species name: *Betabaculovirus spofrugiperdae*; abbreviation: SfGV, (van Oers et al., 2023)) infects the fall armyworm, *Spodoptera frugiperda*, a major pest for maize and other crops (Montezano et al., 2018). Despite its high host specificity —a desirable trait for ecologically friendly pesticides— SfGV has been shown to exhibit slow killing kinetics (Cuartas et al., 2014; Ordóñez-García et al., 2024; Pidre et al., 2019). However, early studies demonstrated that SfGV can enhance nucleopolyhedrovirus (NPV) infection (Shapiro, 2000) and synergize with SfMNPV in certain conditions (Cuartas-Otálora et al., 2019; Ferrelli and Salvador, 2023; Ordóñez-García et al., 2024). To deepen our understanding of the protein components associated with the ODV and the occlusion body matrix of SfGV, we analyzed the protein content of an Argentine isolate of SfGV OBs using MS-based shotgun proteomics. Complemented by bioinformatic analyses of predicted protein structures, we compared our findings with existing proteomic data from other granuloviruses (GVs), to shed light on Betabaculovirus infectious particles.

## Results

### 1. Proteomic profiling of SfGV-ARG occlusion bodies

To initially characterize the purity and morphology of our OB sample, we performed scanning and transmission electron microscopy (**Fig. 1**). Both approaches confirmed the typical ovoidal shape with one single nucleocapsid virion per OB, as reported previously for other granuloviruses (Tsuruta et al., 2018). The OB particle is 442.41 (+/-29,76) nm long and the virion size is 344,34 (+/-29,99) nm (**Fig. 1c**). These sizes are comparable with those reported for the Colombian (SfGV-VG008), SfGV-CH13, and SfGV-CH28 isolates (Cuartas et al., 2014; Ordóñez-García et al., 2024). No evidence of contaminants was detected in our samples, as judged by the homogeneous nature of the OB suspension.

**Figure 1.**
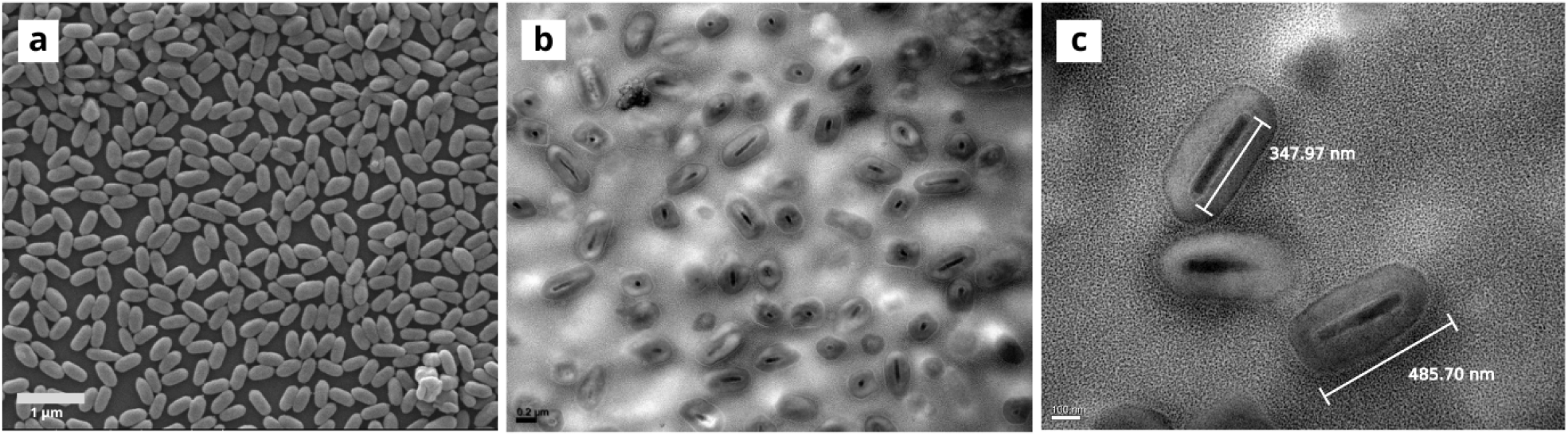
a) Transmission electron microscopy. b) and c) Scanning electron microscopy of SfGV OBs.

After verifying that our purification protocol produces highly homogeneous samples, we cataloged the proteomic composition of SfGV-ARG occlusion particles through liquid chromatography coupled to tandem mass spectrometry (LC-MS/MS). Overall, 72 unique proteins were identified **(Table 1)**, which comprise 47% of the conceptual proteome reported previously for this isolate (Ferrelli et al., 2018). Genes coding for these 72 proteins were distributed all along the genome with no apparent clustering **(Fig. 2)**. Of the 39 core genes reported to date (Cerrudo et al., 2023; Garavaglia et al., 2012; Javed et al., 2017), 29 could be detected in SfGV OB. Ten *per os* infectivity factors, (PIFs 0-9) were described in *Baculoviridae*, all of which are coded in core genes, except for PIF9 (ac108) (Tang et al., 2013). Here, all PIFs were detected except for PIF7 (ORF042) and including ORF045, for which we identified a Baculo_11_kDa (PF06143) domain, that corresponds to PIF9. The remaining 20 core proteins detected in SfGV OB are mainly associated with nucleocapsid and ODV structure. Regarding the 43 non-core proteins detected they include PEP-1, PEP-2, PEP-P10, P10, P24, 38.7k, v-Ubi, Helicase-2, ME53, FP25k, SOD, dUTPase, DBP, P12, PK1, P78/83, ChaB, P13. Of the 7 *bro* genes coded by SfGV, only Bro-F was detected in the OB.

**Table 1.** Proteins detected in SfGV OB.

| ORF | Protein | NCBI ID | AcMNPV homolog | Size (aa) | # Peptides M1 | # Peptides M2 | emPAI M1 | emPAI M2 | emPAI (%VP39) |
| --- | --- | --- | --- | --- | --- | --- | --- | --- | --- |
| 1 | Granulin | AXS01020.1 | Ac8 | 248 | 20 | 26 | 79431,82 | 99999 | 62582,97 |
| 2 | PK1 | AXS01021.1 | Ac10 | 287 | 1 | 13 | 0,16 | 12,34 | 0,97 |
| 3 | P78/83 | AXS01022.1 | Ac9 | 203 | 3 | 3 | 1,68 | 4,18 | 1,86 |
| 5 | P10 | AXS01024.1 | Ac137 | 73 | 4 | 12 | 24,12 | 463,16 | 74,22 |
| 7 | ORF007 | AXS01026.1 |  | 189 | - | 5 | 0 | 2,16 | 0 |
| 9 | ORF009 | AXS01028.1 |  | 190 | - | 7 | 0 | 1,74 | 0 |
| 11 | ODV-E18 | AXS01030.1 | Ac143 | 85 | 6 | 5 | 176,83 | 315,23 | 165,79 |
| 12 | 49k | AXS01031.1 | Ac142 | 454 | 21 | 33 | 5,72 | 99 | 16,71 |
| 14 | PIF5 | AXS01033.1 | Ac148 | 353 | 5 | 15 | 2,16 | 55,23 | 7,67 |
| 16 | PEP1 | AXS01035.1 | Ac131 | 196 | 8 | 11 | 14,85 | 78,43 | 23,96 |
| 17 | PEP2 | AXS01036.1 | Ac131 | 152 | 4 | 12 | 6,5 | 748,89 | 48,99 |
| 18 | PEP/P10 | AXS01037.1 |  | 383 | 5 | 14 | 14,85 | 3980,07 | 170,71 |
| 25 | ORF025 | AXS01044.1 |  | 228 | - | 6 | 0 | 2,51 | 0 |
| 26 | ORF026 | AXS01045.1 |  | 193 | 4 | 7 | 1,15 | 9 | 2,26 |
| 27 | PIF3 | AXS01046.1 | Ac115 | 192 | - | 6 | 0 | 2,16 | 0 |
| 28 | ORF028 | AXS01047.1 |  | 100 | 1 | - | 9 | 0 | 0 |
| 29 | ORF029 | AXS01048.1 |  | 111 | 6 | 4 | 6,5 | 2,16 | 2,63 |
| 36 | p13 | AXS01055.1 |  | 280 | - | 6 | 0 | 1,4 | 0 |
| 38 | PIF2 | AXS01057.1 | Ac22 | 380 | 1 | 11 | 0,12 | 4,62 | 0,53 |
| 40 | ORF040 | AXS01059.1 |  | 1143 | 20 | 62 | 1,13 | 14,85 | 2,88 |
| 43 | v-Ubi | AXS01062.1 | Ac35 | 82 | 2 | 6 | 0,93 | 12,9 | 2,43 |
| 44 | ODV-EC43 | AXS01063.1 | Ac109 | 351 | 11 | 23 | 2,02 | 38,81 | 6,22 |
| 45 | ORF045 | AXS01064.1 | Ac108 | 101 | 1 | 2 | 0,39 | 0,93 | 0,42 |
| 50 | dUTPase | AXS01069.1 |  | 141 | - | 2 | 0 | 0,36 | 0 |
| 52 | SOD | AXS01071.1 | Ac31 | 142 | 3 | 3 | 2,162 | 5,81 | 2,49 |
| 63 | PIF0 | AXS01083.1 | Ac138 | 746 | 2 | 18 | 0,14 | 2,93 | 0,45 |
| 64 | ORF064 | AXS01084.1 |  | 90 | - | 4 | 0 | 4,18 | 0 |
| 68 | P24 | AXS01088.1 | Ac129 | 176 | - | 5 | 0 | 3,64 | 0 |
| 69 | 38.7 k | AXS01089.1 | Ac13 | 173 | - | 2 | 0 | 0,43 | 0 |
| 71 | P10 | AXS01091.1 | Ac137 | 173 | 6 | 12 | 5 | 214,44 | 22,98 |
| 72 | PIF1 | AXS01092.1 | Ac119 | 541 | - | 6 | 0 | 1,4 | 0 |
| 74 | Chtb-2a | AXS01094.1 |  | 162 | 1 | 8 | 0,26 | 6,94 | 0,94 |
| 75 | Chtb-2b | AXS01095.1 |  | 159 | - | 2 | 0 | 0,78 | 0 |
| 77 | DBP | AXS01097.1 | Ac25 | 281 | - | 3 | 0 | 0,64 | 0 |
| 79 | P48/45 | AXS01099.1 | Ac103 | 372 | - | 8 | 0 | 1,61 | 0 |
| 80 | ORF080 | AXS01100.1 |  | 223 | - | 3 | 0 | 0,54 | 0 |
| 81 | P12 | AXS01101.1 | Ac102 | 111 | 1 | 2 | 0,39 | 0,93 | 0,42 |
| 82 | P40 | AXS01102.1 | Ac101 | 372 | 5 | 22 | 0,69 | 22,1 | 2,74 |
| 83 | P6.9 | AXS01103.1 | Ac100 | 58 | 1 | - | 2,16 | 0 | 0 |
| 84 | 38K | AXS01104.1 | Ac98 | 304 | - | 5 | 0 | 1,15 | 0 |
| 87 | PIF4 | AXS01107.1 | Ac96 | 157 | 2 | 4 | 0,59 | 1,51 | 0,66 |
| 88 | ODV-E25 | AXS01108.1 | Ac94 | 219 | 8 | 12 | 20,54 | 176,83 | 42,32 |
| 89 | P33 | AXS01109.1 | Ac92 | 303 | 4 | 9 | 0,69 | 3,81 | 1,14 |
| 90 | P18 | AXS01110.1 | Ac93 | 158 | - | 3 | 0 | 1,15 | 0 |
| 91 | ChaB | AXS01111.1 |  | 79 | - | 2 | 0 | 9 | 0 |
| 96 | VP39 | AXS01116.1 | Ac89 | 324 | 18 | 30 | 16,3 | 1244,2 | 100,01 |
| 97 | ODV-EC27 | AXS01117.1 | Ac144 | 290 | 7 | 13 | 1,45 | 7,8 | 2,36 |
| 98 | BRO-f | AXS01118.1 |  | 370 | - | 12 | 0 | 3,64 | 0 |
| 100 | ORF100 | AXS01120.1 |  | 385 | - | 4 | 0 | 0,62 | 0 |
| 101 | ORF101 | AXS01121.1 |  | 124 | 8 | 8 | 9 | 30,62 | 11,66 |
| 105 | PIF8 | AXS01125.1 | Ac83 | 718 | 7 | 24 | 0,55 | 9 | 1,56 |
| 106 | Ac81 | AXS01126.1 | Ac81 | 191 | 2 | 8 | 0,59 | 9 | 1,61 |
| 107 | GP41 | AXS01127.1 | Ac80 | 294 | 12 | 20 | 17,48 | 214,44 | 42,99 |
| 108 | Ac78 | AXS01128.1 | Ac78 | 103 | 4 | 3 | 13,68 | 20,54 | 11,77 |
| 109 | VLF-1 | AXS01129.1 | Ac77 | 368 | - | 15 | 0 | 4,48 | 0 |
| 112 | Ac75 | AXS01132.1 | Ac75 | 145 | 4 | 11 | 1,31 | 17,74 | 3,38 |
| 114 | Desmoplakin | AXS01134.1 | Ac66 | 637 | - | 5 | 0 | 0,28 | 0 |
| 115 | PIF6 | AXS01135.1 | Ac68 | 139 | - | 6 | 0 | 9 | 0 |
| 120 | FP-25K | AXS01140.1 | Ac61 | 170 | 5 | 9 | 1,42 | 6,02 | 2,06 |
| 125 | Alk-exo | AXS01145.1 | Ac133 | 405 | - | 9 | 0 | 1,42 | 0 |
| 126 | Hel-2 | AXS01146.1 |  | 472 | - | 2 | 0 | 0,17 | 0 |
| 127 | ORF127 | AXS01147.1 |  | 333 | - | 3 | 0 | 0,35 | 0 |
| 130 | ODV-E66 | AXS01150.1 | Ac46 | 674 | 8 | 15 | 1,99 | 14,51 | 3,78 |
| 131 | Enhancin-1 | AXS01151.1 |  | 867 | 14 | 27 | 1,24 | 7,91 | 2,2 |
| 137 | Enhancin-2 | AXS01157.1 |  | 857 | 18 | 37 | 1,83 | 18,31 | 4,07 |
| 140 | Ac53 | AXS01160.1 | Ac53 | 196 | - | 2 | 0 | 0,67 | 0 |
| 141 | ORF141 | AXS01161.1 |  | 417 | 17 | 29 | 6,31 | 135,89 | 20,55 |
| 143 | ORF143 | AXS01163.1 |  | 67 | 3 | 4 | 2,98 | 9 | 3,64 |
| 144 | VP1054 | AXS01164.1 | Ac54 | 324 | - | 9 | 0 | 2,34 | 0 |
| 145 | ORF145 | AXS01165.1 |  | 77 | - | 2 | 0 | 2,16 | 0 |
| 150 | ME53 | AXS01170.1 | Ac139 | 298 | - | 8 | 0 | 1,82 | 0 |
| 151 | ORF151 | AXS01171.1 |  | 99 | - | 2 | 0 | 0,78 | 0 |

**Figure 2.**
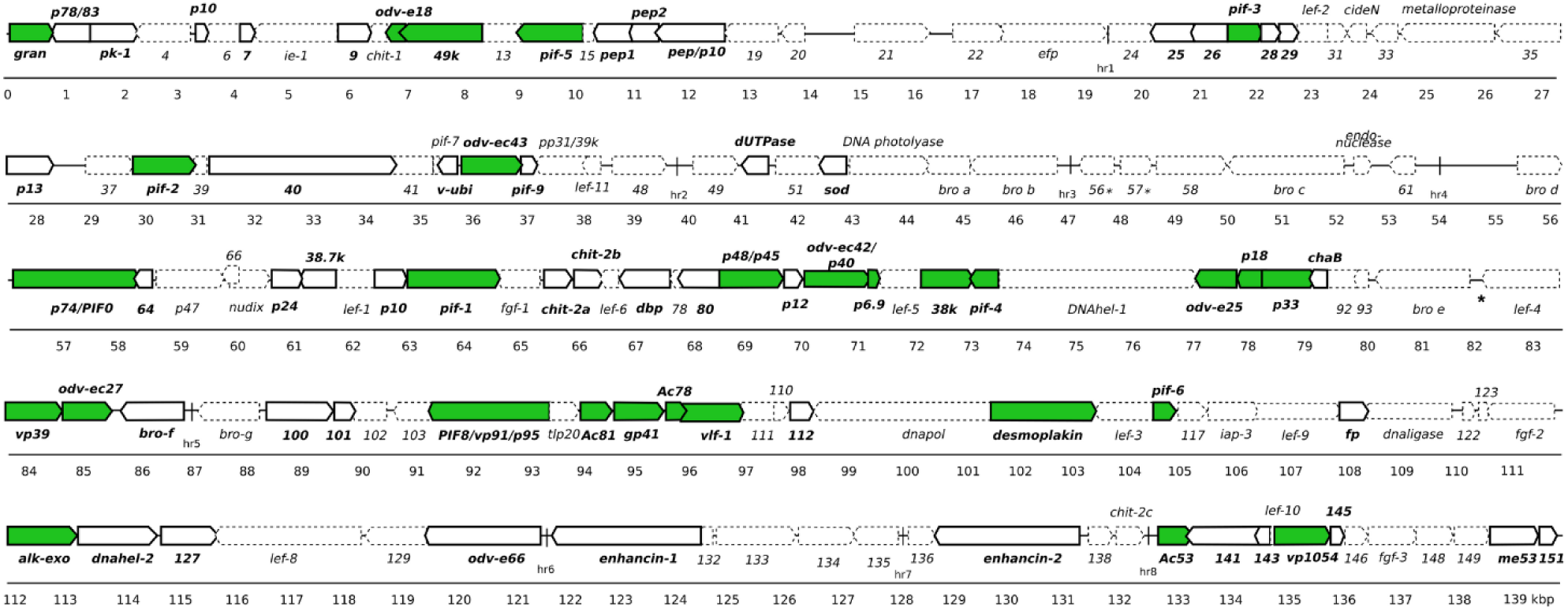
Genomic location of 72 proteins detected in SfGV OB. Protein genes detected in this work are depicted with solid lines. Those corresponding to core genes are shown in green.

Chitin binding proteins were also found in SfGV OB. There are four proteins with predicted chitin binding domains annotated in the SfGV genome, Chit1 (ORF010), Chit2a (ORF074), Chit2b (ORF075), and Chit2c (ORF139). These four proteins are predicted to contain a Carbohydrate-Binding Module family 14 (CBM14), also known as the peritrophin-A domain. In this work, only two of them, Chit2a and Chit2b, were detected in the OB. Although their function is not demonstrated, they likely bind to chitin in the peritrophic membrane, aiding its disruption, in the process of primary infection in the midgut. Bioinformatic and structural analysis of the four chitin binding proteins reveal they contain a putative N-ter transmembrane domain suggesting an ODV envelope anchorage for the two detected. Proteins associated with oral infectivity were also found: ODV-E66 and two enhancins. ODV-E66 is an ODV protein with chondroitinase activity, key for degradation of the peritrophic membrane in primary infection, as demonstrated for its homolog in HearNPV (Hou et al., 2019). On the other hand, enhancin is a metalloprotease coded by several GVs and NPVs that functions as an enhancing factor for viral infection in the insect midgut. It can degrade intestinal mucins present in the peritrophic matrix of Lepidoptera, thereby facilitating the passage of ODV to get into contact with the epithelial midgut cells allowing primary infection (Slavicek, 2012). Two enhancins, Enhancin-1, coded in ORF131, and Enhancin-2, coded in ORF137, were reported in the SfGV genome (Ferrelli et al., 2018) and both were detected in the OB. Sequence and structural analysis show that both enhancins contain an N-terminal peptidase M60 (PF13402) domain and two mucin binding domains in the C-ter domain. This suggests that this protein’s mode of action could involve the binding of the mucin substrate with the C-ter domain and its cleavage with the peptidase, N-ter, domain. Several other proteins found in the OB have no known associated function; these are coded in SfGV ARG orfs 7, 9, 25, 26, 28, 29, 40, 64, 80, 100, 101, 112, 127, 141, 143, 145 and 151. Of particular interest is the detection of ORF040, the largest protein coded in SfGV genome (1143 aa), and ORF028, which is unique to SfGV and has very low sequence complexity. Host proteins were also found in SfGV OB: Ubiquitin-40S ribosomal protein S27a (Uniprot ID: P68203); the cytoskeletal protein Profilin (Uniprot ID: I7G0X7); and the chaperones Heat shock protein 70 A1 (Uniprot ID: A0A0K2CTM7) and Heat shock cognate 70 protein (Uniprot ID:Q8I866). These findings are not surprising, as chaperones, cytoskeleton-related, and ribosomal proteins have previously been reported to be associated with ODVs in other baculoviruses. In particular, Profilin was detected in BmNPV OB, as well as in BV and ODV of MabrNPV and HearNPV. Heat shock cognate 70 protein was found in BmNPV OB, MabrNPV BV and ODV and ClanGV ODV. HSP70 was found in HearNPV BV and ODV, AgMNPV ODV and ClanGV ODV. Additionally, the Ubiquitin-40S ribosomal protein S27a was detected in ClanGV ODV (Braconi et al., 2014; Guo et al., 2017; Hou et al., 2016, 2013; Zhang et al., 2015).

To estimate relative protein levels present in the OB, our analysis included the exponentially modified protein abundance index (emPAI) relativized to the major capsid protein VP39, as reported previously (Masson et al., 2019). Our results show that most of the proteins detected are low in abundance (< 10%). But, considering abundances of >10% empai (%VP39), there are 14 proteins highly abundant in our samples: granulin, pep1, pep2, pep/p10, two P10 homologs (ORF005 and ORF071), six core genes (ODV-E18, VP39, GP41, ODV-E25, 49k, ac78) and two proteins conserved in GVs, with unknown function: ORF101 and ORF141. Bioinformatic analysis of ORF141 showed that it is conserved in GVs and has no conserved domain or similarity with other proteins. Interestingly, homologs of this protein were also detected in EpapGV (Epap126), ClanGV (clan116), and PiraGV (Pr114) proteomic studies (Masson et al., 2019; Wang et al., 2011; Zhang et al., 2015). In EpapGV this protein appeared with mid abundance (Masson et al., 2019). Considering abundances between 1 and 10% there are 16 proteins detected. These include: v-Ubi, SOD, FP-25K, nucleocapsid and ODV specific proteins like PP78/83, Ac81, PIF5, PIF8, P33, Ac75, ODV-EC42/p40, ODV-EC43, and ODV-EC27; and the unknown proteins ORF026, ORF029, ORF040, and ORF143. These unknowns are present in GV genomes, except ORF026 that also have homologs in a few NPVs and asco-and iridovirus. Another interesting finding from the abundance analysis is that Enhancin-2 was nearly twice as abundant as Enhancin-1. As expected, proteins associated with the occlusion matrix or the OB envelope were among the most abundant group (**Fig. 3**).

**Figure 3.**
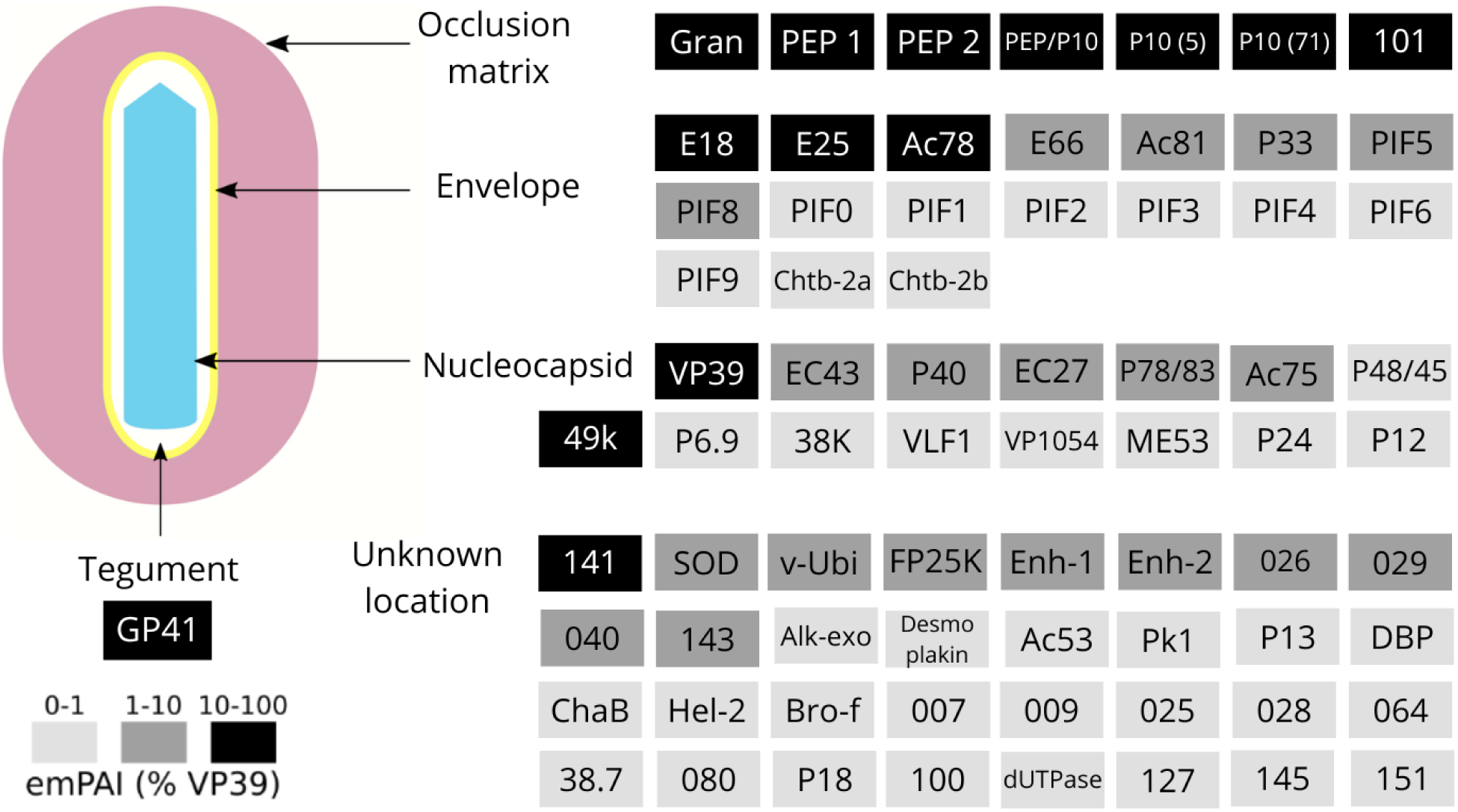
Schematic model of SfGV OB. The 72 detected proteins are grouped according to their predicted location, based on bibliography or bioinformatic analysis. Protein abundance, estimated using the emPAI value, is expressed as a relative value compared to the major capsid protein VP39 and depicted using a grayscale.

### 2. ORF101 is a highly abundant protein and shows similarity to P10 and Ac5

The highly abundant protein coded by orf101 is 124 amino acid long and, interestingly, its homolog in EpapGV (Epap95) was also found highly abundant in the OB (Masson et al., 2019). Homologs were detected in PrGV and ClanGV ODV proteomes, as well (Pr84 and Clan84, respectively). Therefore, to gain insight into the possible function of ORF101 and study its intracellular localization, we expressed a chimeric protein ORF101::GFP in insect cell systems. Interestingly, cells transiently expressing ORF101:GFP protein showed a punctate pattern localized in the cytoplasm, suggesting that this protein can interact with itself forming aggregates (**Fig. 4a**). To better understand these features we analyzed ORF101 bioinformatically. Secondary structure prediction and primary sequence analysis shown in **Fig. 4b** revealed the presence of a N-ter coiled-coil domain, a Proline-rich region, and a basic C-terminal domain with a phosphorylation site (‘KRPS’) overlapping the C-ter basic domain. This site is predicted to be a cAMP-and cGMP-dependent protein kinase phosphorylation site (Prosite, PS00004), whose consensus pattern is [RK](2)-x-[ST] and S or T is the phosphorylation site. The coiled-coil domain supports the possibility of aggregation, as observed in cell culture. However, we also used TAPASS (Falgarone et al., 2022) to see whether it has an amyloid aggregation motif. Prediction of aggregation tendency points to the region 57-71 as the putative amyloid segment causing aggregation. Taken together with the high abundance, aggregation tendency, and sequence organization, we found that ORF101 is very similar to AcMNPV P10.

**Figure 4.**
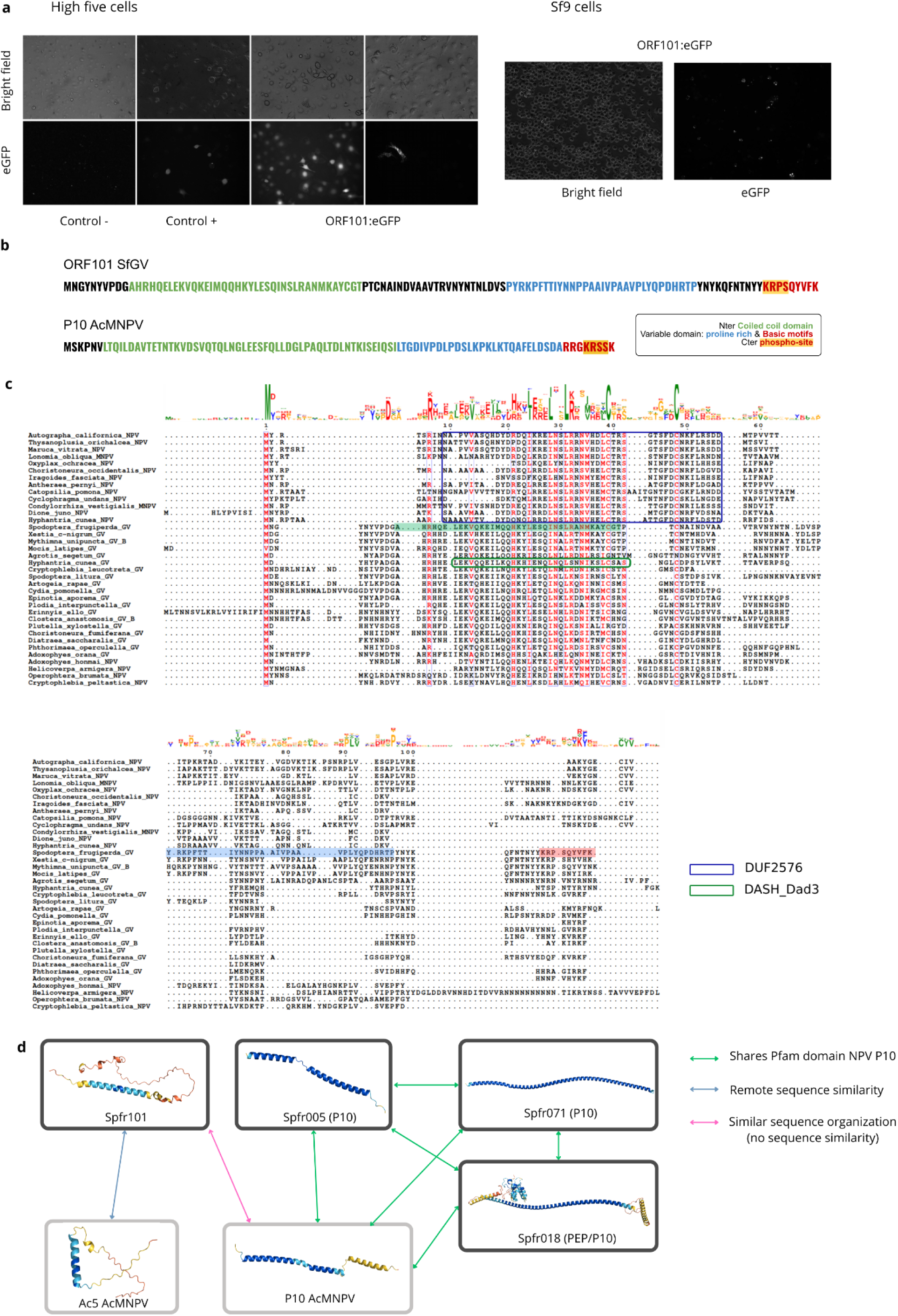
ORF101 analysis. a) Insect cells transfected with ORF101:eGFP constructions. b) ORF 101 and P10 sequence features. c) Multiple sequence alignment of ORF101 and its homologs in GV and NPV. SfGV ORF101 is highlighted in its coiled-coli (green), Proline-rich (blue) and basic motif (red) regions. Pfam domains are squared as indicated in the legend. Sequence IDs are listed in **Table S1.** d) Predicted structures of ORF101 related proteins in SfGV and ACMNPV and their similarities.

P10 is a small protein highly expressed in the very late stage of infection. Studies on AcMNPV P10 showed that this protein forms fibrous structures in the nucleus and in the cytoplasm that are involved in the maturation and release of polyhedra from the nucleus (Carpentier et al., 2008; Graves et al., 2019). P10 associates with microtubules to form these fibrous structures and was shown to interact with host alpha tubulin (Carpentier et al., 2008; Patmanidi et al., 2003). However, it remains unclear whether P10 associates with polyhedra or if it is a structural component of the OB (Carpentier and King, 2009). The P10 sequence is poorly conserved among NPV homologs and consists of an N-ter coiled-coil domain and a C-ter variable domain. The C-ter contains a proline-rich region and a C-ter basic motif (Carpentier et al 2009). The N-ter coiled-coil region is necessary for self aggregation, while the basic C-ter domain is required for the formation of fibrillar structures (van Oers et al., 1993). More recently, it was reported that the C-ter basic domain of AcMNPV P10 is phosphorylated in Ser93 (Raza et al., 2017). This residue is predicted to lie within a cAMP-and cGMP-dependent protein kinase phosphorylation site (Prosite, PS00004). Phosphorylation of P10 regulates its association with microtubules (Cheley et al., 1992; Raza et al., 2017). However, it is unclear which is the microtubule interaction motif in the P10 sequence.

To study ORF101 conservation in the Baculovirus family we first performed a BlastP search that only had hits with hypothetical proteins of betabaculoviruses. All the hits had no recognizable domains except a protein from Hyphantria cunea GV (Uniprot ID: A0A482KC52) which has an N-terminal DASH complex subunit Dad3 domain (PFAM PF08656) overlapping the coiled-coil domain **(Fig. 4c)**. Dad3 is a small but vital subunit of the DASH complex that contributes to the kinetochore-microtubule attachment and ensures accurate chromosome segregation during cell division by coupling microtubule dynamics to chromosome movement (Miranda et al., 2007). This is interesting because this finding could indicate that ORF101 and its GV homologs might interact with microtubules, adding another point of resemblance with AcMNPV P10. The fact that this domain is only detected in one GV (HycuGV), and not in the rest of the homologs, is likely to be due to the highly divergent sequences and the poor representation of baculovirus sequences in the construction of HMM profiles that define domains in the Pfam database.

Then, we performed an hmm-based remote homology search that retrieved NPV homologs revealing that ORF101 is related to the Ac5 protein family (Pfam domain DUF2576; PF10845) **(Table S1)**. Ac5 is found in the AcMNPV OB (Wang et al., 2018), and also its BmNPV homolog (Bm134) was determined to be a good candidate to display fusion proteins in the OB (Guo et al., 2017). Figure 4c shows the conservation of this protein among GVs and NPVs homologs. The multiple alignment shows that the most conserved region overlaps both Pfam domains DUF2576 and DASH complex subunit Dad3, indicating that they might be functionally related. We then predicted the structure of these proteins and found that all of them are simple, non-globular structures made of alpha-helix and unstructured coils, making it difficult to infer homology based on structure similarity. Figure 4d summarizes the relationships among the different proteins mentioned, including 3 additional SfGV proteins that do contain a P10 domain (PF05531).

In conclusion, it appears that ORF101 protein is an OB specific protein of the Ac5 family, with a sequence organization that resembles AcMNPV P10.

### 3. PEP fold is repeated in six OB proteins

To add structural information on the proteins detected in the SfGV OB, we used AlphaFold (Jumper et al., 2021) to predict the tertiary structure of every protein **(Fig. S1)**. With these structures, we aimed to uncover further similarities among OB proteins. Therefore, we used DALI to perform an all-against-all structural comparison to identify similar folds. Interestingly, ORFs 7, 16, 17, 18, 25, and 98 share the same fold **(Fig. 5)**. It is not surprising that similarities were found among pep-1 (ORF 16), pep-2 (ORF 17), and pep/p10 (ORF 18). However, AlphaFold structure predictions of ORFs 7, 25, and 98 showed high structural similarity with the pep fold, as detected by DALI and confirmed by FATCAT structural alignments **(Table S2)**. PEP stands for Polyhedron Envelope Protein, which is known to form a proteinaceous envelope of the OB with several layers (Sajjan and Hinchigeri, 2016). ORF98 codes for Bro-f and it has 3 predicted domains of the PEP fold. To better compare this protein with the others, each domain was aligned separately **(Fig. 5b)**. Psi-blast analysis of ORFs 7, 25 and 98 F found hits only in granuloviruses and there were no crossed hits among them, or with PEPs, meaning that there is not enough sequence similarity among them. Apart from the common fold in all six proteins, there are striking structural similarities between ORF007 and ORF025 but this does not show at the sequence level. These two proteins have unknown functions and Blast searches find homologs only in granuloviruses.

**Figure 5.**
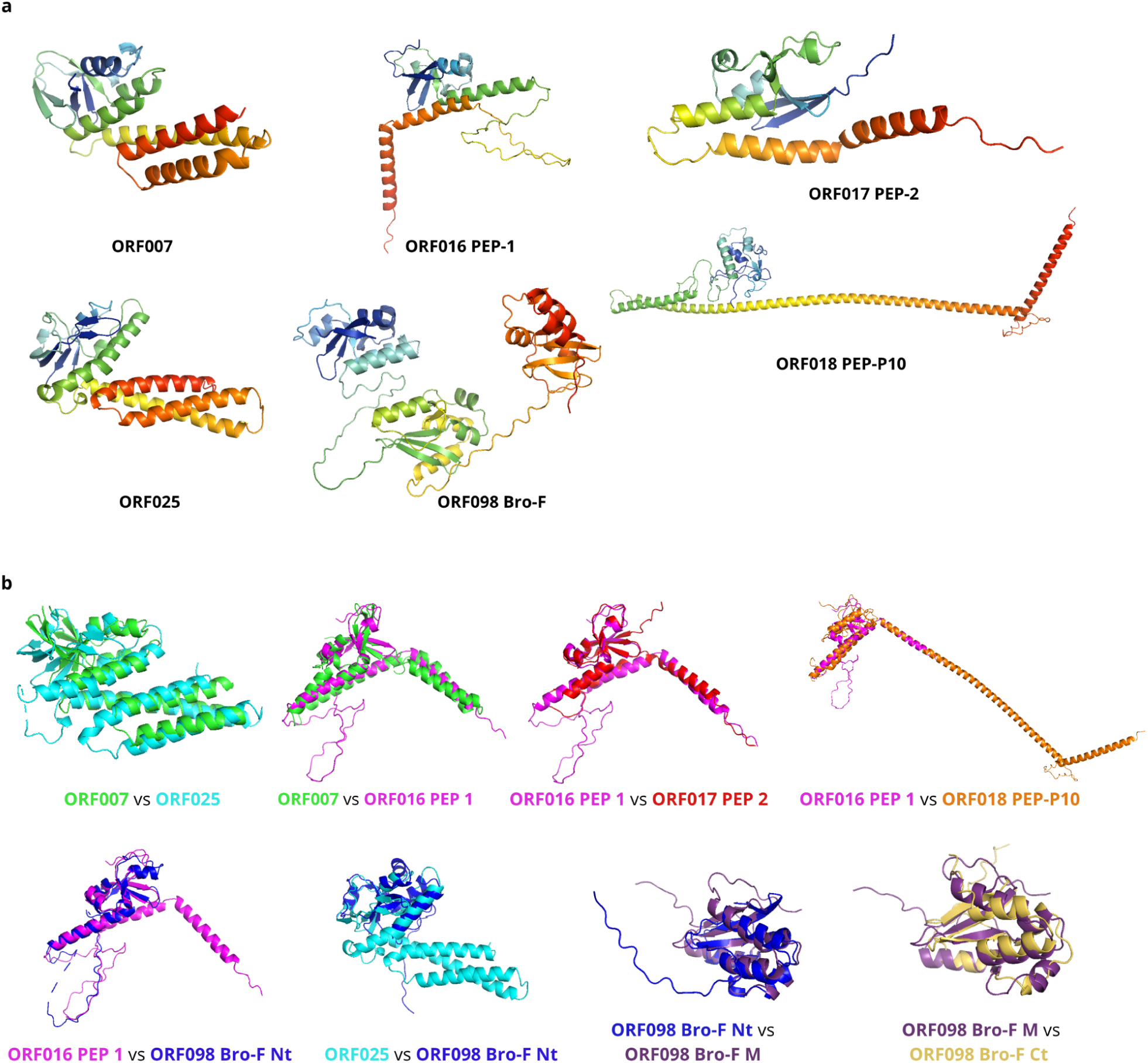
OB proteins that share the PEP fold. a) Structures predicted by Alphafold of ORFs 7, 16, 17, 18, 25 and 98. b) Representative structural alignments by FATCAT of all six proteins. Bro F was chopped into its 3 domains to better show the structural similarity. All the alignments shown are significantly similar. FATCAT alignment scores of all possible combinations are shown in **Table S2**.

**Table S2.** FATCAT flexible structure alignment among PEP similar proteins.

### 4. Proteomic comparison among GVs

To contribute to the general knowledge of betabaculovirus occlusion body composition we compared our data with the available proteomic data for PrGV, ClanGV, and EpapGV (Masson et al., 2019; Wang et al., 2011; Zhang et al., 2015). We used Orthofinder to group all the protein sequences of the four proteomes and, based on the published reports, we determined the common proteins present in the OB (**Fig 6**). We found 33 proteins in common in the four GVs ODV/OB, 23 of which correspond to baculovirus core genes. The 10 remaining proteins include me53, v-ubi, Ac75, bro-f, ORF1629, p12, pep/p10, ORF040, ORF101 and ORF141. Nine proteins were found shared in groups of 3 GVs, which are odv-e25, ac81, pif-3, pk-1, sod, p24, fp, odv-e66, and ORF029, being ac81, pif3 and odv-e25 core genes. Interestingly, 13.7% of the proteins detected in SfGV OB were not shared with other GVs (19 proteins). Regarding the group of proteins detected in the four GVs, there are three proteins with no annotations or conserved domains, i.e. ORF101, ORF040, and ORF141. In SfGV, ORF101 and ORF141 appear as highly abundant proteins. As discussed above, ORF101 is homologous to Ac5 and exhibits striking similarity to P10. We inspected the structures of ORF141 and its homologs and found that the homology is supported by structural similarity (**Fig. S2**). The case of ORF040 is not as straightforward. Although Orthofinder grouped ORF040 with proteins of the other 3 GVs, their sequence and structural similarity are not easily apparent. ORF040 is a 1143 amino acid long protein and their homologs are 803 (PrGVORF42), 802 (Clan22), and 446 (Epap48) amino acid long proteins, respectively. There is considerable variability in their lengths and also in their structures. We could identify that the homologous region is located in the N-ter of the proteins, corresponding to the first 180 amino acids in SfGV ORF040 as shown by multiple sequence alignments (**Fig. S2**).

**Figure 6.**
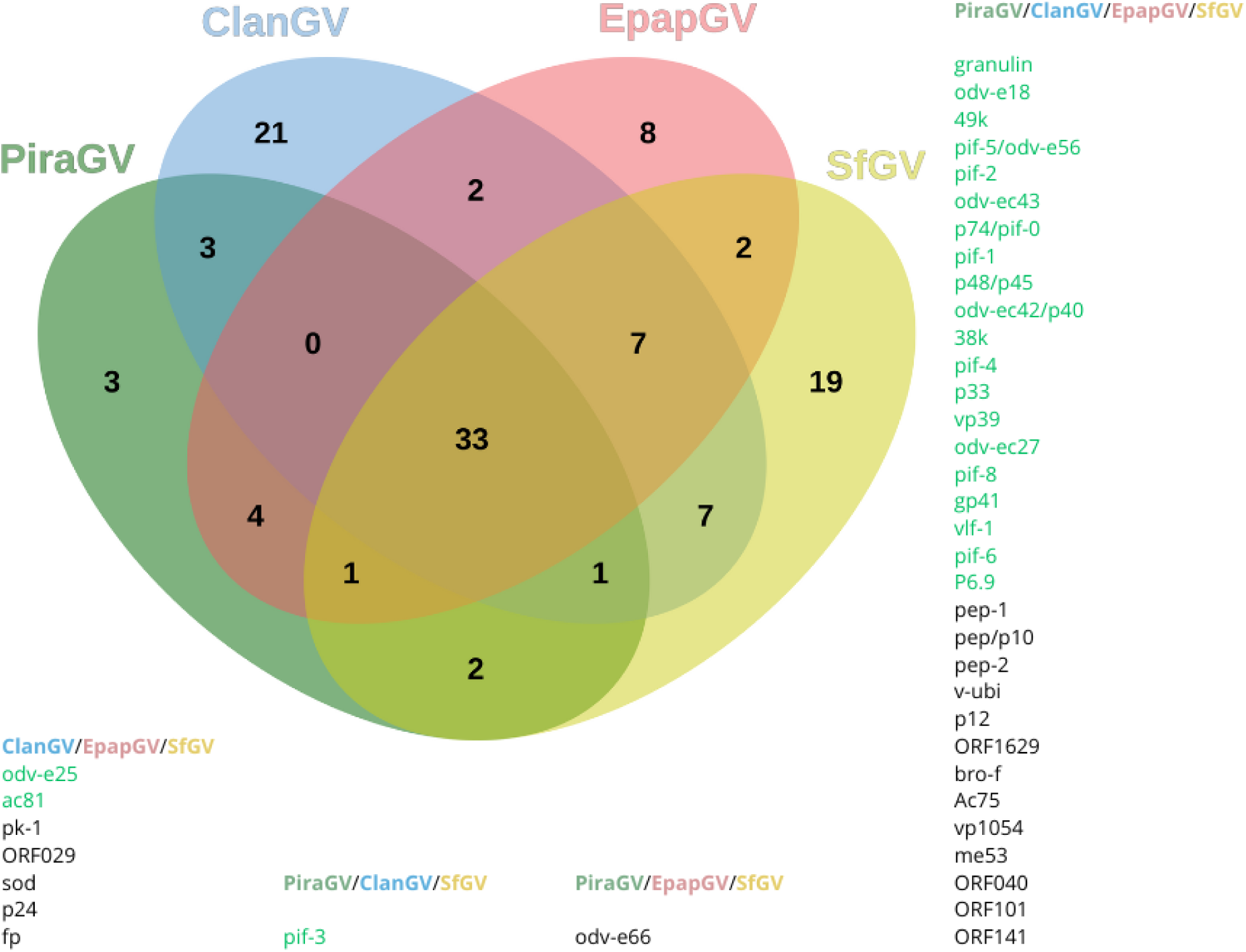
Comparison of GV proteomic studies. The proteins of each intersection are listed in **Table S3**.

## Discussion

Lepidoptera-specific baculoviruses are widely studied due to their importance in pest control. However, Betabaculoviruses are still far behind Alphabaculoviruses in terms of molecular biology studies. Therefore, proteomic analyses provide important insights on the structural components and viral infection. Phylogenies based on core genes divide Betabaculovirus genera in two different clades, “a” and “b” (Miele et al., 2011; Wennmann et al., 2018), where clade “a” contains GVs that infect several insect families and clade “b” is mainly populated with noctuid-specific GVs. In this work we contributed new information on the protein composition of the OB and ODV of SfGV, which belongs to the clade “b”; and compared it with proteomic studies from PiraGV, ClanGV and EpapGV, all belonging to the clade “a”.

It is worth noting that in our work we made no efforts in separating the ODVs from the OB structure before subjecting the samples to proteomics analysis, in a similar way to our previous work about EpapGV (Masson et al., 2019). Conversely, for PiraGV, the authors did separate the fractions, while for ClanGV they used a combination of approaches (Wang et al., 2011; Zhang et al., 2015). Here, according to the proteins commonly detected, we support that the separation of the ODV fraction from the OB matrix does not seem to affect the identification of ODV and OB proteins, allowing a comparison of the four studies.

Most of the proteins detected were as expected, according to current knowledge about nucleocapsid, ODV and OB matrix composition. Recent studies on the structure of the AcMNPV nucleocapsid revealed that it is composed of more than 100 subunits (Jia et al., 2023; Johnstone et al., 2024). The nucleocapsid organization involves a cap, a helical cylinder and a base. It was determined that VP39 (Ac89) is the main component in the helical cylinder and VP80 (Ac104), ODV-EC43 (Ac109), 49k (Ac142), BV/ODV-C42 (P40, Ac101) and ODV-EC27 (Ac144) are components of the head and the base of the nucleocapsid. The base also contains 38K (Ac98) and other proteins not determined in the cryoEM structures but predicted to interact with the base: PP78/83 (ORF1629, Ac9) and P12 (Ac102) (Jia et al., 2023; Johnstone et al., 2024). All these proteins were detected in common in the 4 GV studies **(Fig. 6)**, with the exception of VP80. This is in accordance with the fact that VP80 is conserved in lepidopteran NPV genomes and absent in GVs. Other nucleocapsid-associated proteins are Ac75, P48 (P45, Ac103), VP1054, ME53 and P24, and were also detected in SfGV and the other GVs, as well.

ODV associated proteins were also expected. *Per os* infectivity factors (PIFs) are essential for infection of midgut cells. PIF proteins assemble in a complex anchored in the ODV membrane and mediate interaction with epithelial cell receptors. There are ten PIF proteins, termed PIF-0 to PIF-9, coded in core genes, with the exception of PIF-9 (Ac108) which is found in all alpha, beta, and gamma baculoviruses (Garavaglia et al., 2012). All of them were found in this study, except for PIF-7, a 53 aa protein coded in orf042. All were shared in 4 GVs compared, only PIF-3 was not detected in EpapGV. This supports a common way of primary infection in all baculoviruses, considering that this complex was only studied in NPVs and not in GVs (Boogaard et al., 2018).

Apart from the PIFs, other ODV proteins were found as well, such as ODV-E66, which was considered a putative PIF protein (Boogard 2018), and the baculovirus core proteins ODV-E18, ODV-E25, Ac78, Ac81 and P33. In summary, all core genes that code for structural components of the nucleocapsid, ODV or OB matrix were detected in SfGV; whereas core proteins related to RNA or DNA polymerase were not present in the OB, supporting the reliability of the proteomics technique used.

Alphafold has revolutionized the protein structure prediction field (Jumper et al., 2021). In this work we used Alphafold2 to predict every SfGV protein structure to deepen our understanding of granulovirus virion and OB composition. We found a common fold among six proteins in the OB, that strikingly matches the N-ter globular domain present in PEP. Due to the known function associated with PEP in forming the envelope of the baculovirus OB, and adding the fact that these 6 proteins were detected in the SfGV OB, it is tempting to hypothesize that all of them might be implicated in a similar structural role in the envelope. The case of **Bro-f** is also striking due to the fact that there are several *bro* genes coded in SfGV but only this one was present in the OB. Whatsmore, the function of bro genes is not well defined. In addition, when comparing proteomes of the 4 GV analyzed so far, we detected that Bro-f is present in all of them, as well as Pep-1 pep-2 and pep-p10. ORF007 is present only in SfGV OB and ORF025 was also found in ClanGV (Clan32).

The MS-based proteomics technique used, allowed us to perform protein semi-quantification based on the emPAI value (Ishihama et al., 2005). This, in turn, permitted highlighting the importance of some proteins according to their abundances. One of the most abundant proteins found in SfGV OB is ORF101, which is also highly abundant in EpapGV (Masson et al., 2019), and has been detected in ClanGV and PiraGV proteomes. We found that it belongs to the Ac5 family, which has been described as a polyhedron protein in AcMNPV, with potential biotechnological applications (Guo et al., 2017; Wang et al., 2018). It is striking the similarity found in the sequence organization of ORF101 and P10, indicating these proteins could have similar functions. However, we could not detect homology between ORF101 and P10 by means of sequence-based remote homology search. We inspected their predicted structures, but they are too simple to assure homology based on structure similarity. Conversely, we were able to detect remote homology to Ac5, inferring that ORF101 could be an occlusion matrix protein. Overall, the alpha helix predicted structures of SfGV proteins ORF101, ORF005, ORF071 and ORF018 and AcMNPV proteins P10 and Ac5 suggest that these proteins could interact with others forming fibrils, as was demonstrated for P10 (Carpentier et al., 2008; Graves et al., 2019).

Another highly abundant protein in SfGV is ORF141 which was interestingly found in the four GV proteomes. Its homolog in EpapGV, Epap126, was found with intermediate abundance in EpapGV OB (Masson et al., 2019). It is worth noting that the analysis of abundance and the comparative proteomics allowed highlighting ORF101 and OFR141 as possibly important proteins in granuloviruses, and therefore as candidate proteins for further study.

Granuloviruses are not as well studied as NPVs, so protein localization in the OB is primarily inferred by the knowledge of their NPV orthologues. Therefore we infer that pep-1, pep-2 and pep-p10 are part of the OB envelope, in a similar way that NPV PEP does. Similarly, we consider the P10 homologs ORF005 and ORF071 as part of the OB **(Fig. 3)**, mainly due to their high abundance. However, although NPV P10 is not associated with the virion structure, it is not confirmed if it is actually part of the polyhedron in AcMNPV. Finally, we also assume ORF101 is part of the occlusion matrix, due to the similarity found with Ac5 and its high abundance.

Interestingly, there is a set of 19 proteins detected in SfGV that is not shared with other GVs. ORF028 is included, which is the only SfGV specific protein, with no homologs in Genbank, so confirming that this hypothetical protein is actually expressed. This group also includes three core genes, Ac78, desmoplakin and ac53, but their detection only in SfGV and not in the other 3 GVs, indicates their role in the virus structure is probably not essential. On the other hand, enhancins are coded in SfGV but not in EpapGV, ClanGV or PiraGV. Enhancins are metalloproteinases capable of degrading insect intestinal mucin, facilitating the passage of ODVs through the peritrophic matrix and reaching midgut cells to establish primary infection (Wang and Granados, 1997). SfGV was reported to have an enhancing activity and this is associated with the enhancin protein in the granule (Shapiro, 2000). SfGV ARG was reported to code for two enhancin proteins (Ferrelli et al., 2018) and here we confirmed the presence of both in the OB. Bioinformatic and structural analysis of both proteins reveal their structures are highly similar, even though they share only 22.7 % identity (not shown). Also, apart from the typical peptidase domain already known for enhancins, here we could also detect mucin binding motifs in the C-terminal region in accordance to a recent report (Ricarte-Bermejo et al., 2021).

Understanding baculovirus OB structure is an important field of investigation due to their importance in primary infection and in its growing biotechnological applications (Fabre et al., 2020; López et al., 2024). The detection of all proteins in SfGV OB provides insights into SfGV and granulovirus OB composition, paving the way for a deeper understanding of the roles of OB proteins in the bioinsecticide applications of baculoviruses.

## Materials and Methods

### Larvae and virus

*Spodoptera frugiperda* larvae were reared at the Institute of Microbiology and Agricultural Zoology (IMYZA), INTA, and IBBM. Larvae were raised on a maize-based artificial diet (Greene et al., 1976) at 25 ± 1 °C, 16:8 h L:D, 50–70% RH. An Argentinian isolate of Spodoptera frugiperda granulovirus, SfGV ARG (Pidre et al., 2019), was used for OB protein detection and its genome for peptide mapping (GenBank: MH170055.1).

### Occlusion bodies (OBs) preparation

Larvae were infected with SfGV OBs as described previously (Ferrelli et al., 2018). Dead larvae, with typical signs of infection, were collected and homogenized in double-distilled water (ddH2O), gauze filtered, and resuspended in 0,1% sodium dodecyl sulfate. After 2 min of low speed centrifugation (1000 x) to eliminate cell debris, the supernatant with viral occlusion bodies (OBs) was centrifuged at 12000 x g. Pellets containing OBs were washed twice and finally resuspended in ddH2O. Then, OBs were loaded onto a discontinuous 35–60% (w/w) sucrose gradient and centrifuged at 20000 rpm and 4° C for 1 h in a SW41 rotor (Beckman). The whitish/opalescent band containing OBs was removed, diluted and washed twice in ddH2O. Finally, OBs were centrifuged at 14000 x g for 10 min and resuspended in ddH2O.

### Scanning and Transmission Electron Microscopy (SEM and TEM)

For SEM, an OBs sample was fixed with 4 % paraformaldehyde in PBS at 4°C for 24 h. Then, OBs were centrifuged for 5 min and 8000 x g and resuspended in 100 % ethanol, and subjected to critical point drying and gold sputtering. Image acquisition was performed in an environmental scanning electron microscope (ESEM, FEI, model Quanta 200). For TEM, a diluted, translucid OB sample was incubated for 2 h with 1 % paraformaldehyde in PBS. Then, OBs were centrifuged for 5 min and 8000 x g. This fixation process was repeated twice but using 2% paraformaldehyde in PBS. A secondary fixation was performed with 1 % osmium tetroxide for 1 hour at 4°C. Then, samples were dehydrated using a series of increasing alcohol concentrations and finally included in an epoxy resin. Ultrathin sections were cut and then contrasted with uranyl acetate and lead citrate and examined under a transmission electron microscope JEM 1200 EX II (JEOL Ltd., Tokio, Japan).

### Sample processing and mass spectrometry analysis

Two biological independent samples were processed. The total protein mass in the sample was quantified using the Bradford assay (Bradford, 1976). OB samples were treated with SDS-PAGE sample buffer (SDS 0.1 % p/v and β-mercaptoethanol) for 5 min at 100 °C and then loaded on a 12% p/v polyacrylamide running gel combined with a 4% stacking gel and electrophoresed for 30 min and 120 V. After Coomassie staining, protein bands were excised from the gel and fractionated in 4 slices for further MS processing.

Samples were sent to the Proteomics Core Facility CEQUIBIEM, at the University of Buenos Aires/CONICET (National Research Council) for protein digestion and analysis. Protein samples were reduced with 10 mM dithiothreitol in 50 mM ammonium bicarbonate pH 8 (45 min, 56°C) and carbamidomethylated with 20 mM iodoacetamide in the same solvent for 40 min, room temperature and in darkness. Then 0.2 volumes of 100% w/v trichloroacetic acid (Sigma) was added, incubated at -20°C for at least two hours and centrifuged at 12000 x g for 10 min (4°C). The pellet was washed twice with ice-cold acetone and dried at room temperature. Proteins were resuspended in 50 mM ammonium bicarbonate (pH 8) and digested with trypsin (Promega V5111). The resulting peptides were desalted with ZipTip C18 columns (Millipore).

The digested sample was analyzed by nanoLC-MS/MS in a Thermo Scientific Q Exactive Mass Spectrometer coupled to a nano HPLC EASY-nLC 1000 (Thermo Scientific). For the LC-MS/MS analysis, the column was loaded with 1 μg of peptides and eluted for 120 min using a reverse phase column (C18, 2 μm x 10 nm particle size, 50 μm x 150 mm) Easy-Spray Column PepMap RSLC (P/N ES801) which separates complex mixtures of peptides with a high degree of resolution. A flow rate of 300 nL min^-1^ was used for the nano-column. The solvent range was from 7% B (5 min) to 35% B (120 min). Solvent A was 0.1% formic acid in water and B was 0.1% formic acid in acetonitrile. The injection volume was 2 μl. The MS equipment has a high collision dissociation cell (HCD) for fragmentation and an Orbitrap analyzer (Thermo Scientific, Q-Exactive). For ElectroSpray Ionization (Thermo Scientific, EASY-SPRAY) a voltage of 3.5 kV was used.

XCalibur 3.0.63 (Thermo Scientific) software was used for data acquisition and equipment configuration to identify peptides simultaneously with their chromatographic separation. Full-scan mass spectra were acquired in the Orbitrap analyzer. The scanned mass range was 400–1800 m/z, at a resolution of 70000 at 400 m/z and the 12 most intense ions in each cycle were sequentially isolated, fragmented by HCD and measured in the Orbitrap analyzer. Peptides with a charge of +1 or with an unassigned charge state were excluded from fragmentation for MS2.

Q Exactive raw data was processed using Proteome Discoverer^TM^ software (version 2.1.1.21, Thermo Scientific) and searched against SfGV ARG protein database (GenBank: MH170055.1) digested *in silico* with trypsin with a maximum of one missed cleavage per peptide. Proteome Discoverer^TM^ searches were performed with a precursor mass tolerance of 10 ppm and a product ion tolerance of 0.05 Da. Static modifications were set to carbamidomethylation of Cys, and dynamic modifications were set to oxidation of Met and N-terminal acetylation. Protein hits were filtered for high confidence peptide matches with a maximum protein and peptide false discovery rate of 1% calculated using a reverse database strategy. The exponentially modified protein abundance index (emPAI) was calculated automatically by Proteome Discoverer^TM^ software and used to estimate the relative abundance of identified proteins within the sample. The emPAI value for the major capsid protein VP39 was used to normalize protein abundance. A geometric mean EmPAI in the two replicates was calculated for each protein and this was then normalized to the same mean for VP39. The abundance of each protein was then expressed as %VP39 and classified as low (<1%VP39), middle (1 - 10% VP39), and high (>10% VP39).

### ORF101-GFP construction and visualization

SfGV orf101 was amplified from SfGV-ARG DNA by PCR using primers Fw (5’ CTGCAA<u>GGTACC</u>CCAATTATTTGCGAGACCAA3’) and Rev (5’ CTGCAA<u>GAGCTC</u>TTATTTAAACACATATTGAC 3’) containing recognition sites for KpnI and SacI (underlined). The PCR product was cloned between KpnI and SacI sites in the plasmid pIP-eGFP (Fabre et al., 2020) to obtain a orf101:eGFP construct under the control of the OpIE2 promoter from Orgyia pseudotsugata MNPV for constitutive expression of the fusion protein. As a positive control, we used an in-house GFP expression plasmid. As a negative control we constructed pIP-eGFP unable to express eGFP, by digesting the plasmid with KpnI and SacI and re-ligating the plasmid, thereby eliminating eGFP start codon.

HighFive and Sf9 insect cell lines were grown at 27 °C in Grace’s (InvitrogenTM) medium containing 10 % fetal bovine serum (FBS, Internegocios S.A., Argentina). The recombinant plasmid was transfected in both insect cell lines using Cellfectin (Thermo) following manufacturer’s instructions. Briefly, 8 x 10^5^ cells resuspended in Grace’s medium with 1.5 % FBS were pleated in wells from 6-well plates. Eight μl of Cellfectin were diluted in 100 μl of medium and 1 μl of plasmid was diluted in 100 μl of medium. Then, both dilutions were mixed together and incubated for 15 min at room temperature and poured onto the cells. Transfected cells were incubated at 27°C and monitored under fluorescent microscopy for 5 days.

### Bioinformatic analysis of ORF101

ORF101 protein sequence was subjected to Quick2D tool in MPI toolkit site and Prosite for determination of coiled-coil regions and phosphorylation sites, respectively (Gabler et al., 2020; Sigrist et al., 2013). TAPASS server was used to identify putative aggregation sites (Falgarone et al., 2022). Homology search was performed with BlastP (NCBI) and Jackhmmer (Altschul et al., 1990; Potter et al., 2018). The latter was used to find remote homologs of ORF101 in Alphabaculovirus. Representative sequences of ORF101 and Ac5 homologs were aligned with mafft (Katoh et al., 2002). The multiple sequence alignment was visualized in www.alignmentviewer.org, which outputs the accompanying sequence logo, shown in Fig 4c.

### Protein structure prediction and similarity search

Protein structure prediction was performed with Alphafold2 algorithm (Jumper et al., 2021) through the Colabfold notebook (Mirdita et al., 2022) for all 72 SfGV proteins detected in the proteomics assay. We subjected 72 SfGV structures to the DALI all-against-all comparison tool (Holm et al., 2023). The resulting similar structures were aligned with Flexible structure AlignmenT by Chaining Aligned fragment pairs allowing Twists (FATCAT) (Li et al., 2020), to confirm similarities. The PyMOL Molecular Graphics System, Version 1.74 Schrödinger, LLC. was used for structure visualization and manipulation, and for figure production.

### Comparative proteomics analysis

To analyze the shared proteins in the four GVs with reported proteomics studies, we first determined orthologous proteins among them. The complete conceptual proteomes of ClanGV (HQ116624.1), PiraGV (GQ884143), EpapGV (NC_018875.1) and SfGV (MH170055.1) were downloaded from NCBI and were subjected to Orthofinder v2.5.5. (Emms and Kelly, 2019). Orthogroups were further processed with in-house python scripts to add bibliographic data on presence/absence of each protein in their respective proteomic study, unify protein names, and obtain comparable lists of detected proteins for each virus. These lists were intersected in jvenn (Bardou et al., 2014) to obtain the Venn diagram. Webserver or local Hmmscan (v 3.3) was performed with all the proteome of SfGV against Pfam database, to detect non annotated domains. Also, predicted structures were submitted to Foldseek (Kempen et al., 2022) to confirm domain similarities by structural comparison, when necessary.

## Acknowledgements

This research was funded by grants from Agencia Nacional de Promoción Científica y Tecnológica (ANPCyT): PICT 2017–0758 and PICT 2020 SERIE A 02466 to M.L. Ferrelli and PICT 2014–1827 to V. Romanowski.

## Supplementary material

**Figure S1.** Protein structure prediction of 72 proteins detected in SfGV OB.

**Figure S2.** Structural similarity of ORF040 and ORF141 homologs.

**Table S1.** Uniprot IDs of proteins used in Figure 4c MSA.

**Table S2.** FATCAT flexible structure alignment among PEP similar proteins.

**Table S3.** Shared proteins in four GVs.

