## Supplementary material for "Proteomic Characterization of Spodoptera frugiperda Granulovirus Occlusion Bodies": Figs S1 and S2

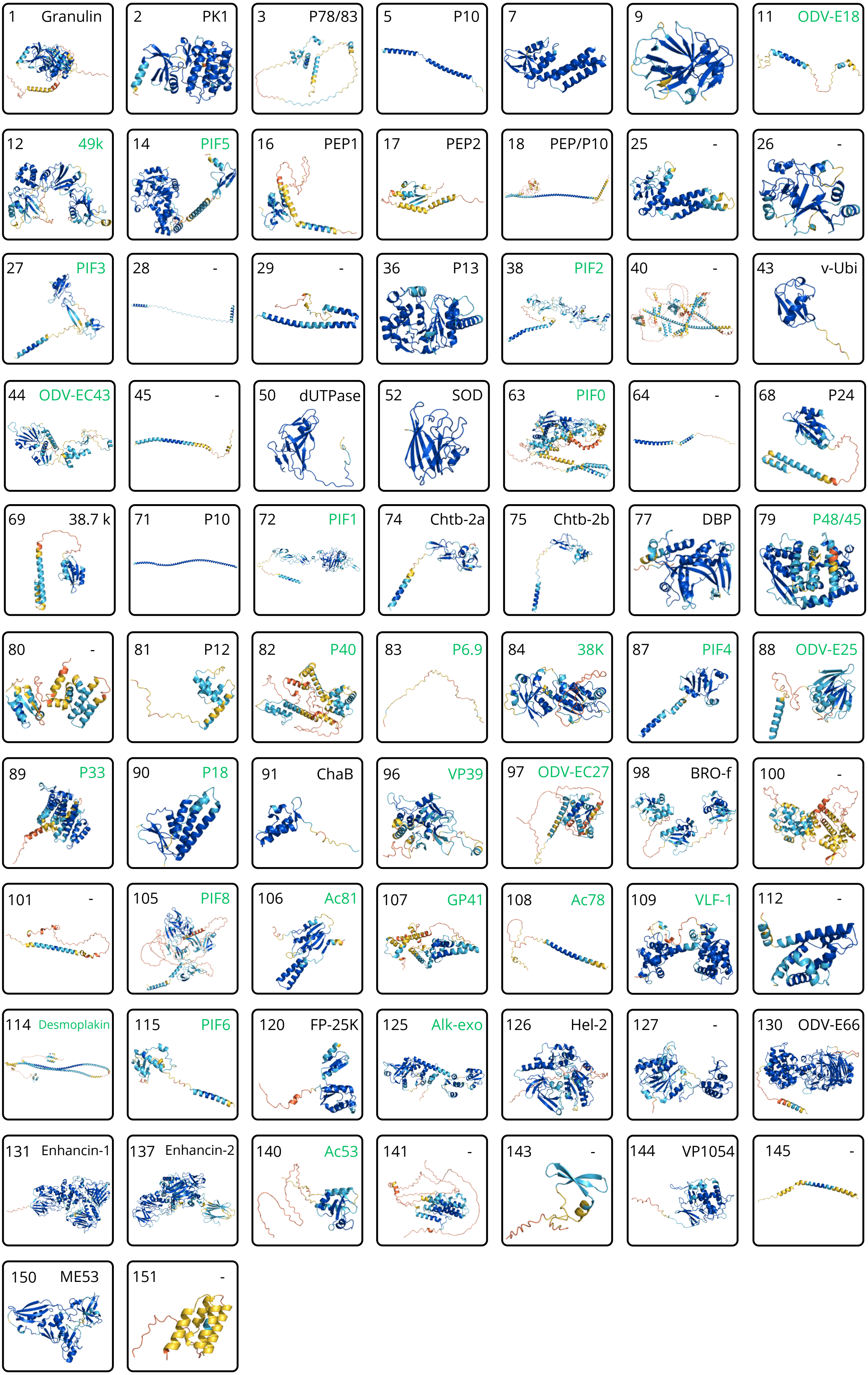
1 Granulin
2 PK1
3 P78/83
5 P10
7
9
11 ODV-E18
12 49k
14 PIF5
16 PEP1
17 PEP2
18 PEP/P10
25 -
26 -
27 PIF3
28 -
29 -
36 P13
38 PIF2
40 -
43 v-Ubi
44 ODV-EC43
45 -
50 dUTPase
52 SOD
63 PIF0
64 -
68 P24
69 38.7 k
71 P10
72 PIF1
74 Chtb-2a
75 Chtb-2b
77 DBP
79 P48/45
80 -
81 P12
82 P40
83 P6.9
84 38K
87 PIF4
88 ODV-E25
89 P33
90 P18
91 ChaB
96 VP39
97 ODV-EC27
98 BRO-f
100 -
101 -
105 PIF8
106 Ac81
107 GP41
108 Ac78
109 VLF-1
112 -
114 Desmoplakin
115 PIF6
120 FP-25K
125 Alk-exo
126 Hel-2
127 -
130 ODV-E66
131 Enhancin-1
137 Enhancin-2
140 Ac53
141 -
143 -
144 VP1054
145 -
150 ME53
151 -

Figure S2. Protein structures of ORF141 and ORF040 and their homologues.

**ORF141**

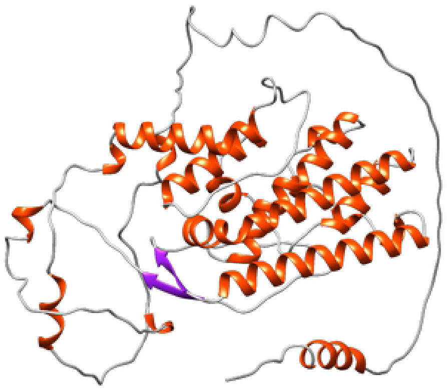

SfGV (ORF141)

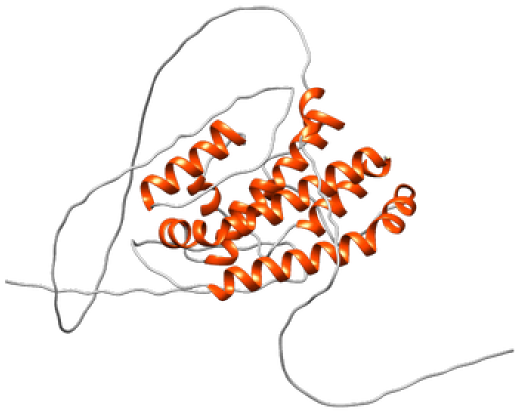

PiraGV ()

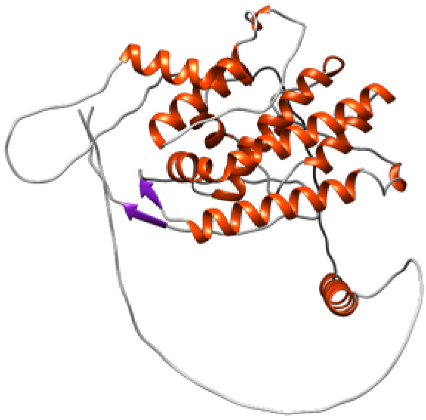

EpapGV ()

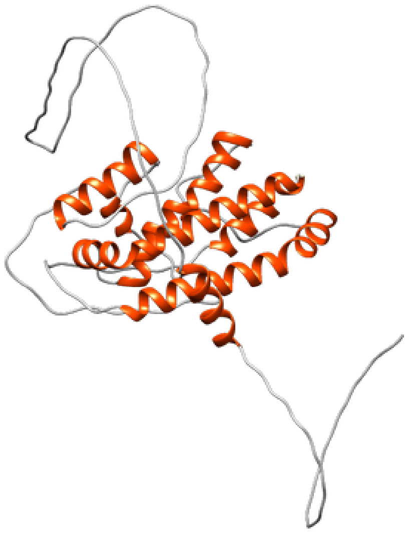

ClanGV ()

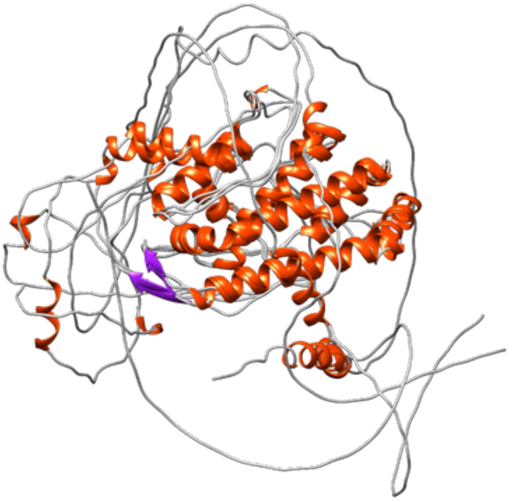

ORF141 multiple alignment

**ORF040**

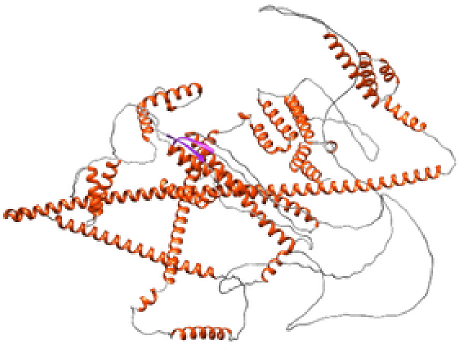

SfGV (ORF040)

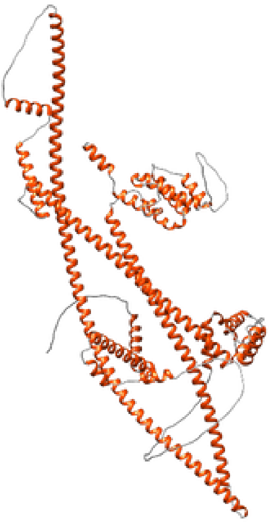

PiraGV (PrGVORF42)

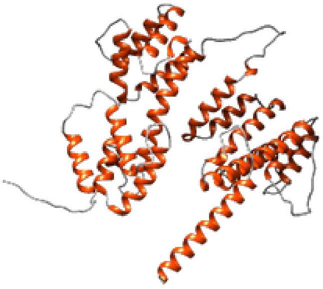

EpapGV (Epap48)

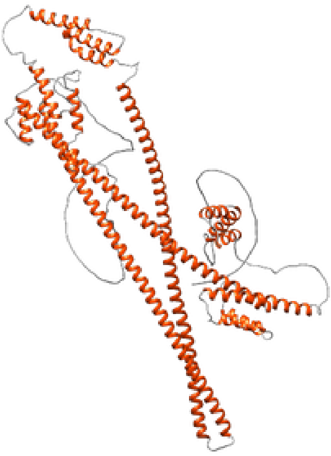

ClanGV (Clan22)

### ORF040

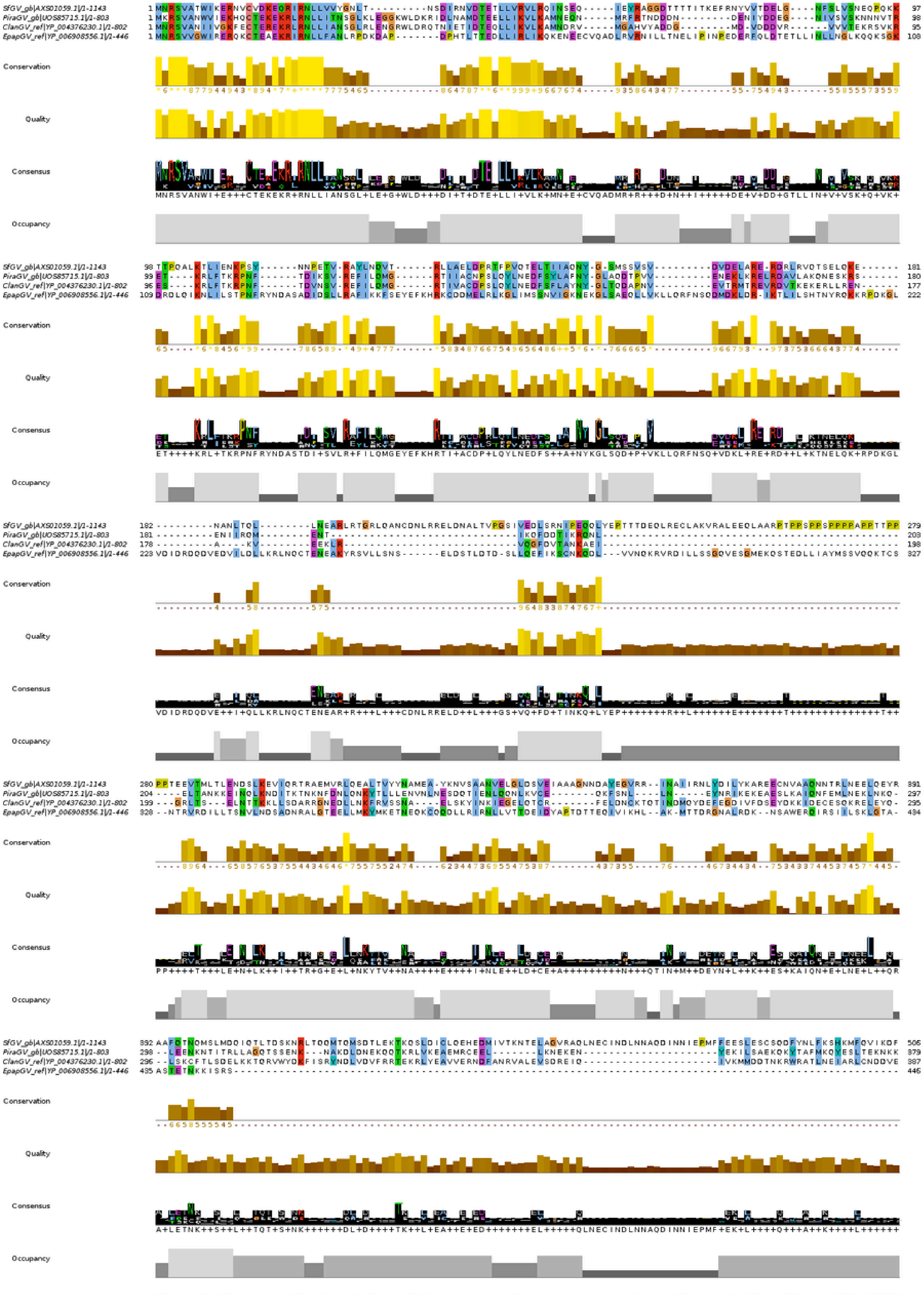
SfGV_gb|AXS01059.1|/1-1143
PiraGV_gb|UOS85715.1|/1-803
ClanGV_ref|YP_004376230.1|/1-802
EpapGV_ref|YP_006908556.1|/1-446
Conservation
Quality
Consensus
Occupancy

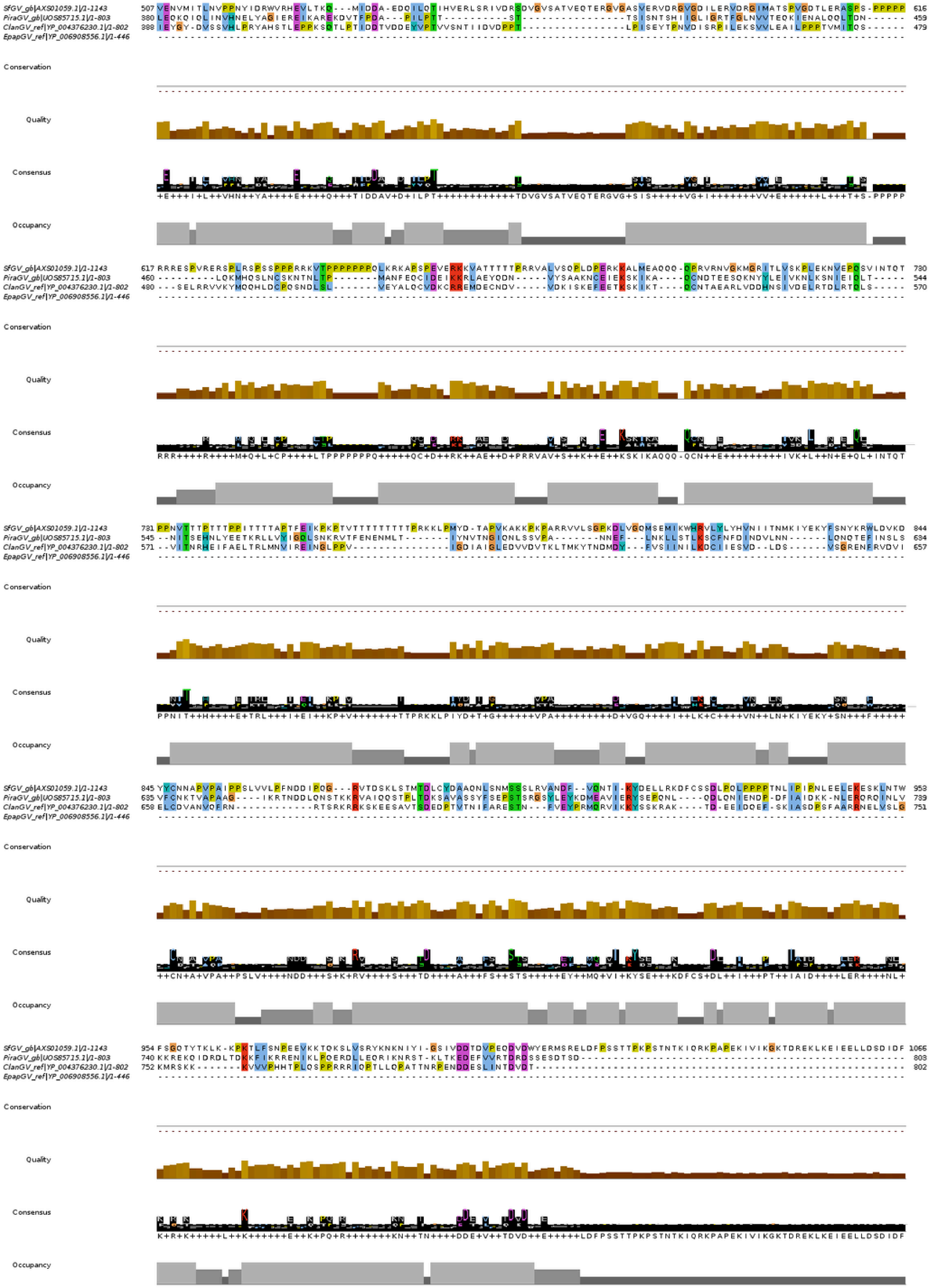

SfGV_gb|AXS01059.1|/1-1143
PiraGV_gb|UOS85715.1|/1-803
ClanGV_ref|YP_004376230.1|/1-802
EpapGV_ref|YP_006908556.1|/1-446
Conservation
Quality
Consensus
Occupancy

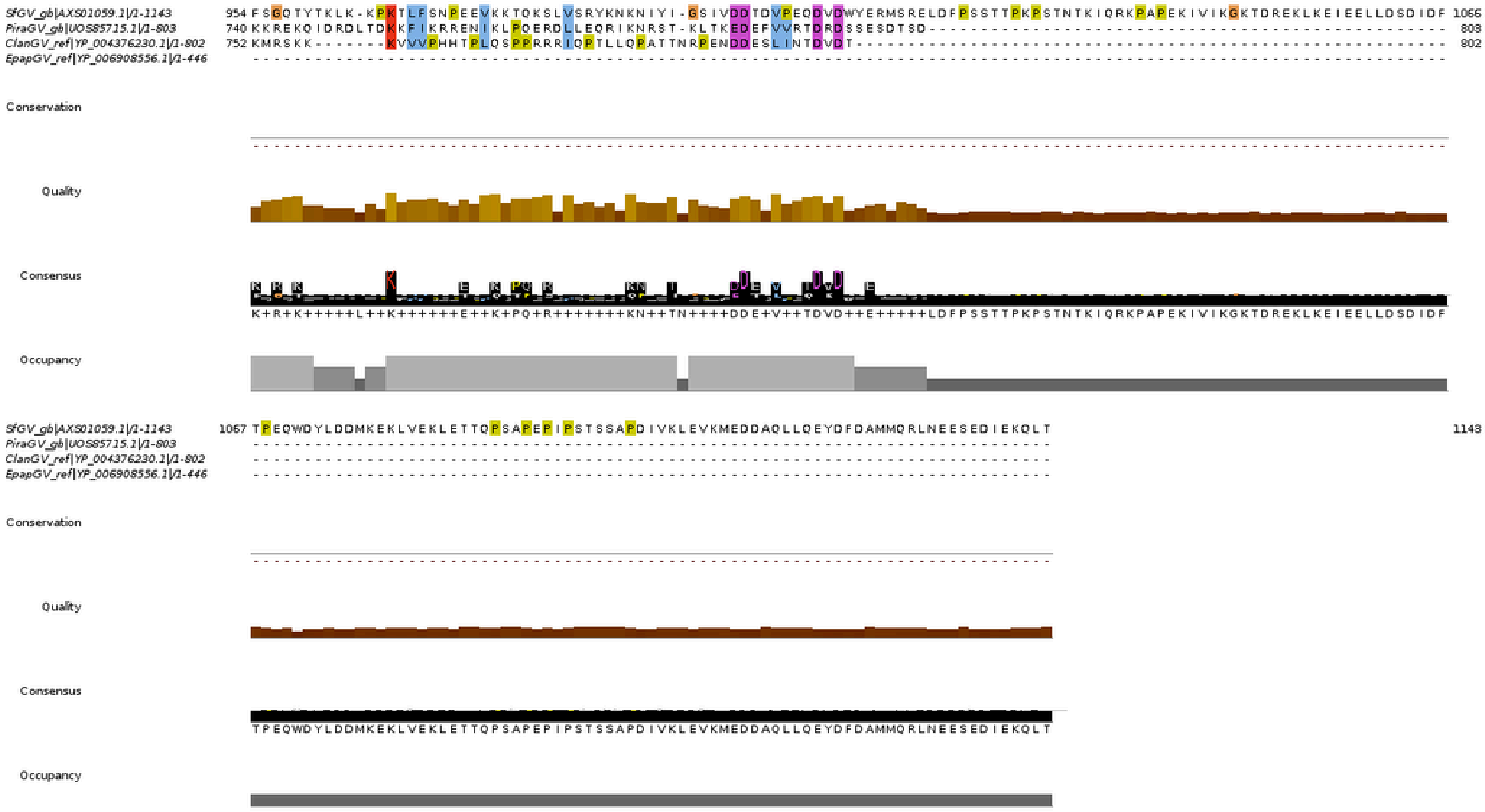
SfGV_gb|AXS01059.1|/1-1143 954 FSGQTYTKLK-KPKTLFSNPEEVKKTQKSLVSRYKNKNIYI-GSIVDDTDVPEQDVDWYERMSRELDFPSSTTPKPSTNTKIQRKPAPEKIVIKGKTDREKLKEIEELLDSDIDF 1066
PiraGV_gb|UOS85715.1|/1-803 740 KKREKQIDRDLTDKKFIKRRENIKLPQERDLLEQRIKNRST-KLTKEDEFVVRTDRDSSESDTSD-------------------------------------------------- 803
ClanGV_ref|YP_004376230.1|/1-802 752 KMRSKK--------KVVVPHHTPLQSPPRRRIQPTLLQPATTNRPENDDESLINTDVDT-------------------------------------------------------- 802
EpapGV_ref|YP_006908556.1|/1-446
Conservation
Quality
Consensus
K+R+K+++++L++K++++++E++K+PQ+R++++++++KN++TN++++DDE+V++TDVD++E+++++LDFPSSTTPKPSTNTKIQRKPAPEKIVIKGKTDREKLKEIEELLDSDIDF
Occupancy
SfGV_gb|AXS01059.1|/1-1143 1067 TPEQWDYLDDMKEKLVEKLETTQPSAPEPIPSTSSAPDIVKLEVKMEDDAQLLQEYDFDAMMQRLNEESEDIEKQLT 1143
PiraGV_gb|UOS85715.1|/1-803
ClanGV_ref|YP_004376230.1|/1-802
EpapGV_ref|YP_006908556.1|/1-446
Conservation
Quality
Consensus
TPEQWDYLDDMKEKLVEKLETTQPSAPEPIPSTSSAPDIVKLEVKMEDDAQLLQEYDFDAMMQRLNEESEDIEKQLT
Occupancy
